# Transcriptomic Stratification of Late-Onset Alzheimer’s Cases Reveals Novel Genetic Modifiers of Disease Pathology

**DOI:** 10.1101/763516

**Authors:** Nikhil Milind, Christoph Preuss, Annat Haber, Guru Ananda, Shubhabrata Mukherjee, Cai John, Sarah Shapley, Anna L. Tyler, Benjamin A. Logsdon, Paul K. Crane, Gregory W. Carter

## Abstract

Late-Onset Alzheimer’s disease (LOAD) is a common, complex genetic disorder well-known for its heterogeneous pathology. The genetic heterogeneity underlying common complex diseases poses a major challenge for targeted therapies and the identification of novel disease-associated variants. Case-control approaches are often limited to examining a specific outcome in a group of heterogenous patients with different clinical characteristics. Here, we developed a novel approach to define relevant transcriptomic endophenotypes and stratify decedents based on molecular profiles in three independent human LOAD cohorts. By integrating post-mortem brain gene co-expression data from 2114 human samples with LOAD, we developed a novel quantitative, composite phenotype that can better account for the heterogeneity in genetic architecture underlying the disease. We used iterative weighted gene co-expression network analysis (WGCNA) analysis to reduce data dimensionality and to isolate gene sets that are highly co-expressed within disease subtypes and represent specific molecular pathways. We then performed single variant association testing using whole genome-sequencing data for the novel composite phenotype in order to identify genetic loci that contribute to disease heterogeneity. Distinct LOAD subtypes were identified for all three study cohorts (two in ROSMAP, three in Mayo Clinic, two in Mount Sinai Brain Bank). Single variant association analysis identified a genome-wide significant variant in *TMEM106B* (p-value < 5×10^−8^, rs1990620^G^) in the ROSMAP cohort that confers protection from the inflammatory LOAD subtype. Taken together, our novel approach can be used to stratify LOAD into distinct molecular subtypes based on affected disease pathways.

## INTRODUCTION

Late-onset Alzheimer’s disease (LOAD) is the most common form of dementia in the elderly. The clinical features associated with LOAD are an amnesic type of memory impairment, deterioration of language, and visuospatial deficits. In the later stages of the disease, symptoms may include motor and sensory abnormalities, gait disturbances, and seizures. Without advances in therapy, the number of symptomatic cases in the United States is predicted to rise to 13.2 million by 2050^1^.

Many common, complex diseases such as LOAD present with heterogeneous phenotypes due to interactions between genetic and environmental factors affecting a range of pathways and processes. LOAD has no simple form of inheritance and is governed by a common set of risk alleles across multiple genes that, in combination, have a substantial effect on disease predisposition and age of onset^2^. Genome-Wide Association Studies (GWAS) have become an important tool for identifying variants in complex diseases^3,4^. GWAS for LOAD have identified variants in over 500 genes as potential risk factors with the ε4 variant in *APOE* as the strongest contributor to overall disease risk^2,5^. LOAD has a strong polygenic component and an estimated heritability of up to 80%^6^. It has been challenging to transition from the identification of associated genetic variants to the molecular mechanisms that lead to the accumulation of amyloid plaques and helical tau filaments^7^. Furthermore, there is mounting evidence that the observed heterogeneity in LOAD is associated with multiple distinct subtypes^8,9^.

Gene co-expression modules tend to consist of genes that belong to the same cellular pathways or programs and help explain the global properties of the transcriptome as it relates to disease risk^10^. Networks-based co-expression module approaches have been used to identify causal variants in Late-Onset Alzheimer’s disease^7,11^. However, such studies have failed to account for the heterogeneity of mechanisms that lead to complex diseases. Here, we analyze whole genome sequencing (WGS) and whole transcriptome data from three independent human cohorts from the Accelerating Medicines Partnership - Alzheimer’s Disease (AMP-AD) Consortium. We use gene co-expression modules to develop quantitative phenotypes that account for the complex genetic architecture and heterogeneity of LOAD to more effectively map associated variants using genome-wide assocation. Furthermore, the method presented in this paper can be used to identify variants in other complex diseases.

## METHODS

### Whole genome sequencing and RNA sequencing data

We obtained whole-genome sequencing and RNA sequencing (RNA-Seq) data from Synapse (https://www.synapse.org/) for three cohorts from the AMP-AD consortium, from the Mayo Clinic, Mount Sinai Brain Bank, and Rush University. The Mayo Clinic (Mayo) cohort consists of 276 temporal cortex (TCX) samples from 312 North American Caucasian subjects consisting of cases characterized with LOAD, pathological aging (PA), progressive supranuclear palsy (PSP), or elderly controls^12^ (Synapse:syn5550404). The Mount Sinai Brain Bank (MSBB) cohort consists of 214 frontopolar prefrontal cortex (FP), 187 inferior temporal gyrus (IFG), 160 parahippocampal gyrus (PHG), and 187 superior temporal gyrus (STG) samples characterized with LOAD, elderly control, or mild cognitive impairment (MCI) (Synapse: syn3159438). The Rush University’s Religious Orders Study and Memory and Aging Project (ROSMAP) cohort consists of 623 dorsolateral prefrontal cortex (DLPFC) samples of individuals from 40 groups of religious orders from across the United States (ROS) and older adults in retirement communities in the Chicago area (MAP), characterized with LOAD, elderly control, or MCI^7,13^ (Synapse:syn3219045). A summary of samples from each of the cohorts is provided in Table S1 and Table S2. Sex, age of death, and batch were used as covariates for normalization in the ROSMAP and Mayo data. Sex, age of death, race, and batch were used as covariates for normalization in the MSBB data. Details on post-mortem brain sample collection, tissue and RNA preparation, sequencing, and sample quality control can be found in published work related to each cohort^12,14,15^.

### Co-expression modules and iterativeWGCNA

Data on human AMP-AD co-expression modules were obtained from Synapse (Synapse: syn11932957.1). The modules derive from the three independent LOAD cohorts used in this study. A detailed description on how co-expression modules were identified can be found in a recent study that identified the human co-expression modules as part of a transcriptome wide LOAD meta-analysis^16^. In brief, a modified procedure using five different co-expression analysis protocols followed by merging by graph clustering methods was performed to obtain 30 modules across all three cohorts (Synapse: syn2580853), 26 of which corresponded to the six tissue regions used in this study. A summary of these modules is provided in Table S3. We focused on tissues from the frontal cortex, temporal cortex, and hippocampus due to their relevance to LOAD neuropathology^17^. These modules are generally large, containing thousands of genes that represent multiple functions^16^. In order to construct more functionally-specific submodules from these AMP-AD co-expression modules, we subjected them to a repeated pruning process called iterativeWGCNA^18^. Briefly, iterativeWGCNA performed WGCNA on each AMP-AD co-expression module independently. The gene sets produced by this process were then pruned to ensure that only highly correlated genes remained by evaluating the connectivity of the genes to the gene set eigengene. The resulting gene sets, containing highly correlated genes, were combined and the process was repeated until the gene sets converged. The algorithm then attempted to reclassify genes from the residual gene set. We specified a soft-threshold power of six, a minimum eigengene connectivity of 0.6, and a required module size of 100 to promote the generation of submodules that capture pathway-level signals. The final set of 68 submodules consisted of highly correlated and cell-type specific genes. The submodules were mutually exclusive for a given cohort but overlapped with submodules from other cohorts. A summary of these submodules is provided in Table S4. An eigengene for a given submodule is defined as the first principle component of gene expression data within each submodule.

### Stratification of LOAD cases based on clustering of human co-expression submodules

Eigengene expression data for TCX, PHG, FP, and DLPFC regions was used to stratify LOAD cases in separate analyses. Clustering was performed on submodule eigengenes to determine subtypes of LOAD cases in each brain region. The NbClust R package determined the optimal number of clusters for different clustering methods by polling with the majority rule across 30 indices^19^. We tested agglomerative hierarchical approaches (Ward, UPGMA, WPGMA) and a reallocation approach (K-means) on the eigengene expression data and evaluated the within-cluster similarity of cases using silhouettes. The silhouette for a given object is a measure that simultaneously assesses how similar the object is to its cluster and how different the object is from all the other clusters^20^. Prior analysis of simulated genome-wide methylation data suggests that no one clustering method outperforms the other consistently and that mean silhouette widths can be used to pick the ideal clustering method^21^. The silhouette plots revealed that different methods were required for the different regions to generate clusters with the largest average silhouette widths. We determined that K-means was an optimal approach for DLPFC, Ward was optimal for PHG and TCX, and UPGMA was optimal for FP after analyzing silhouette plots of clusters generated by each method for each region. An example of silhouettes used to determine the ideal clustering method for the DLPFC region is shown in Figure S1. A summary of the clusters for each brain region, considered case subtypes, is provided in Table S5. In the subtypes generated for the DLPFC region from the ROSMAP cohort, we assessed each subtype for enrichment of cognitive and pathological measures. We used Braak stages as a measure of neurofibrillary tangle burden and CERAD scores as a measure of neuritic plaque burden^22,23^. We also assessed the rate of decline in memory, executive function, visuospatial function, and language across the subtypes. Definitions, collection, and standardization of these decline measures can be found in previously published work^24^.

### Differential expression analysis of case subtypes

For differential expression analysis, control decedents were defined as cognitively-normal and MCI decedents for PHG, FP, and DLPFC. In the case of TCX, control decedents were defined as cognitively normal, PSP, and PA decedents. For each of the regions used to stratify LOAD cases (TCX, PHG, FP, and DLPFC), we performed differential expression analysis to compare gene expression in case subtypes with control decedents as described above. We used the limma R package to perform the differential expression analysis between subtype and control decedents^25^. We used the clusterProfiler R package to perform KEGG and Reactome pathway analysis on differentially expressed genes to determine the signal captured by clustering on eigengene expression data^26^.

### Single-variant association of eigengene expression and subtype specificity

We used EMMAX, a variance component linear mixed model, to perform single-variant association of our newly derived quantitative traits^27^. Each submodule eigengene was used as a quantitative trait in single-variant association for its respective brain region. For each region, we also developed a subtype specificity metric by calculating the Euclidean distance between the eigengene expression profile of each decedent and the centroid of each subtype cluster. This resulted in a vector of scores for each subtype that was mapped separately. All quantitative trait mapping results had a genomic inflation factor near one, indicating that there was no significant population substructure effect on the mapping. QQ plot analysis on the p-values showed no evidence of population substructure or confounding effects (Figure S2).

### Replication of suggestive and significant SNPs in other cohorts

The ROSMAP cohort represented the most adequately powered cohort in the study and was used as a baseline for assessing replication of suggestive and significant SNPs in the other cohorts. SNPs were considered suggestive if quantitative trait mapping with either the submodule eigengenes or the subtype specificity metric resulted in a p-value smaller than 1×10^−5^ and genome-wide significant if they resulted in a p-value smaller than 5×10^−8^, which are standard cutoffs for GWAS. Suggestive and significant SNPs from the DLPFC region in ROSMAP were considered replicated in the TCX, FP, and PHG regions if the SNPs were associated with the submodule eigengenes or subtype specificity metric of the given region at a p-value of 0.05. Summary statistics of prior association studies were obtained from the NHGRI-EBI catalog^28^. Loci were considered replicated if suggestive and significant SNPs from the ROSMAP cohort were reported in these studies at a p-value smaller than 5×10^−8^. A summary of the entire analysis is provided in Figure S3.

## RESULTS

### Refinement of 26 human co-expression modules identifies disease-associated transcriptomic signals

We performed an iterative gene list pruning process using the iterativeWGCNA approach to refine the 26 human co-expression modules from the AMP-AD consortium. This resulted in subsets, or submodules, of highly correlated genes that were exclusive to each module. Genes that were not highly correlated to any submodule were removed since they are less likely to contribute to the overall signal of the submodule and more likely to introduce noise. We compared the submodules and detected specific LOAD-associated molecular pathways and processes that are shared across the three post-mortem brain cohorts and six brain regions (Figure S4). Furthermore, incorporating information from previously defined cell-type specific markers derived from bulk RNA-Seq and single cell RNA-Seq^29^ showed that pruning the 26 co-expression modules into 68 submodules resulted in multiple novel cell-type specific submodules (Figure 2, Figure S5). Taken together, these novel 68 submodules reflect 15 specific functional consensus clusters that are associated with distinct pathways and processes related to LOAD (Figure S4).

**Figure 1:**
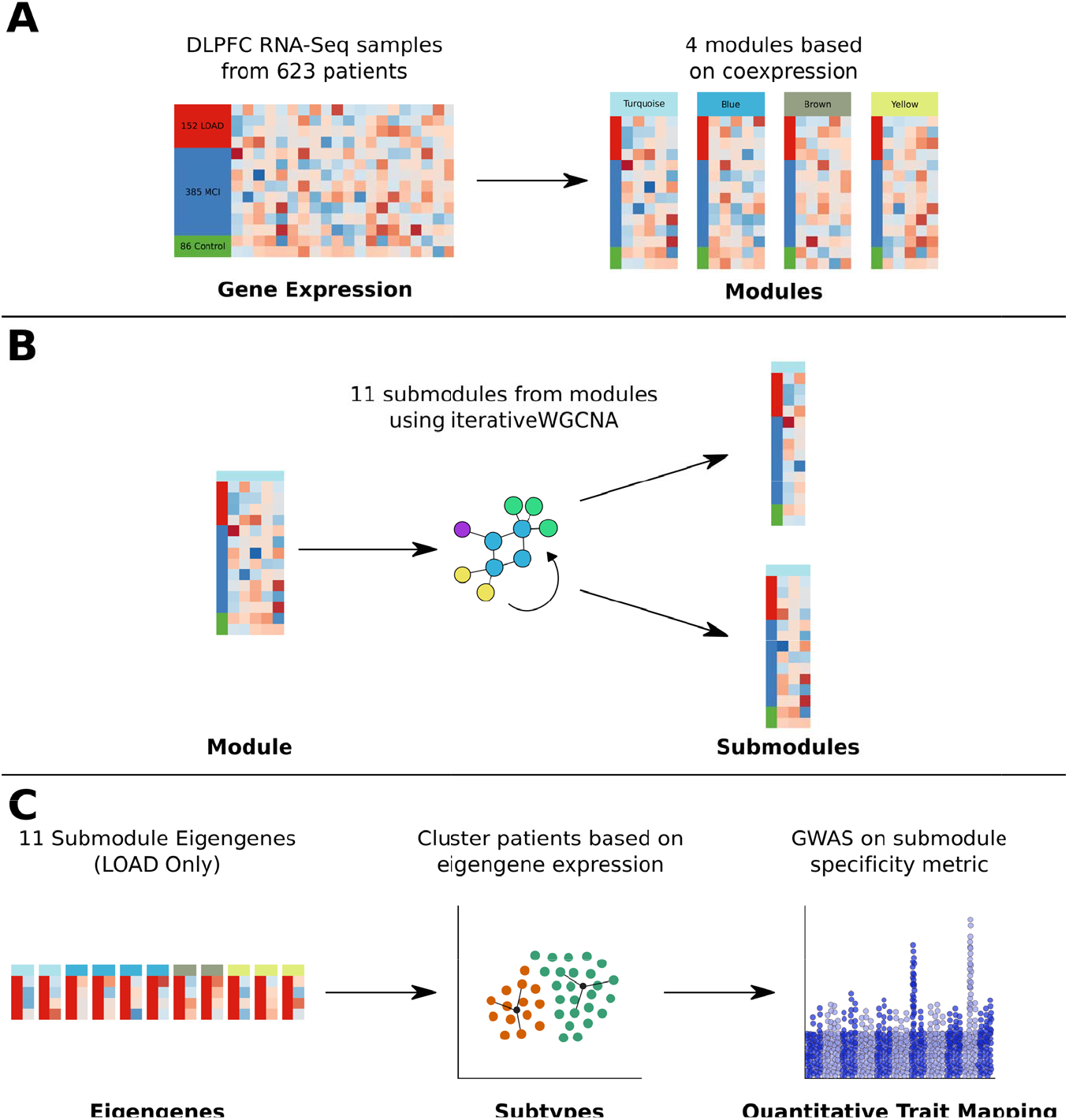
Methodology used in this study to map genetic drivers of LOAD pathology in the ROSMAP cohort. (A) RNA-Seq was performed on dorsolateral prefrontal cortex (DLPFC) tissue from 86 control decedents, 385 decedents with Mild Cognitive Impairment (MCI), and 152 decedents with Late-Onset Alzheimer’s Disease (LOAD). A modified procedure using seven different WGCNA protocols, followed by merging by clustering methods, was performed to obtain 4 modules based on gene co-expression. (B) Each of the four modules was subjected to iterativeWGCNA, a procedure that repeatedly performs WGCNA on expression data to generate highly correlated gene sets and exclude weakly correlated genes. 11 submodules were generated from the 4 modules. (C) The eigengene, or first principal component of each submodule, was calculated for all 11 submodules and used as a quantitative trait for single-variant association mapping. Furthermore, the eigengene expression for LOAD cases was used to perform cluster analysis and generate subtypes of LOAD cases. A Euclidean distance quantitative trait was developed to identify genomic loci for each subtype using single-variant association mapping.

**Figure 2:**
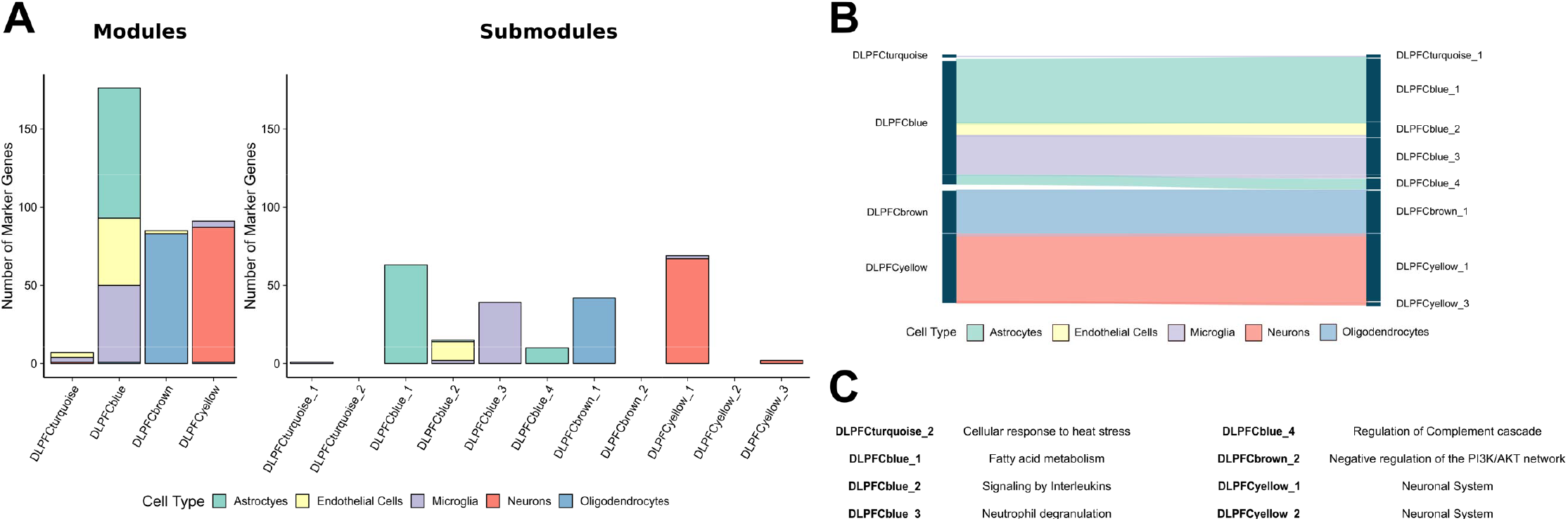
Cell-type specificity of modules is refined in submodules. (A) Cell type specific marker genes reported by McKenzie *et al.* were used to annotate modules and submodules for astrocytes, endothelial cells, microglia, neurons, and oligodendrocytes. The top 100 marker genes for each cell type were used. The iterativeWGCNA procedure generated submodules that were more cell-type specific than their modules of origin. (B) A Sankey diagram demonstrates which cell-type specific markers from modules were found in submodules generated using iterativeWGCNA for the ROSMAP cohort. (C) Gene set enrichment analysis for Reactome pathways was performed for each submodule gene list. The top enriched Reactome pathways for submodules are reported.

### Submodule gene sets capture biological signals specific to LOAD pathology

We annotated submodules using GO term enrichment, KEGG pathway enrichment, and Reactome pathway enrichment to highlight the biological specificity of co-expression signals captured by the different submodules (Table S6, Table S7, Table S8). While the 26 harmonized co-expression modules were associated with five distinct consensus clusters that captured a broader signal, the submodule associations were more specific in terms of functional enrichment (Figure S4). The 15 functional consensus clusters associated with the 68 submodules revealed cell-type specific signatures and elucidated gene sets for specific biological pathways, including tau-protein kinase activity, neuroinflammation, myelination, and cytoskeletal reorganization (Figure S4).

### Single-variant association mapping of submodule eigengenes

To map the genetic drivers of biological disease-associated signals resolved by submodules, we performed single-variant association mapping of submodule eigengenes. Eigengenes were defined as the first principle component of the gene expression data associated with each submodule. They capture the variation of gene co-expression and reduce noise associated with the transcriptomic data. Genome-wide suggestive and significant loci were detected for submodules in all four brain regions (Table S9, Table S10, Table S11, Table S12). We identified multiple loci that were replicated across the cohorts at a genome-wide significant level. For instance, rs1990620 is a known variant in *TMEM106B* that was identified as genome-wide significant in the DLPFC region from the ROSMAP cohort was replicated (p < 5×10^−2^) in all other brain regions from the Mayo and MSSM cohorts.

### Stratification of LOAD cases based on 68 AMP-AD co-expression submodules

Clustering LOAD cases in subtypes based on eigengenes provided a method of assessing genetic drivers of heterogeneity in the transcriptome of LOAD cases. The NbClust package chose between two and three clusters for each region and the number of cases in each cluster was balanced (Table S5). The subtypes were not enriched for common LOAD-associated covariates, such as sex, *APOEε4* genotype, or years of education (Figure 4).

**Figure 3:**
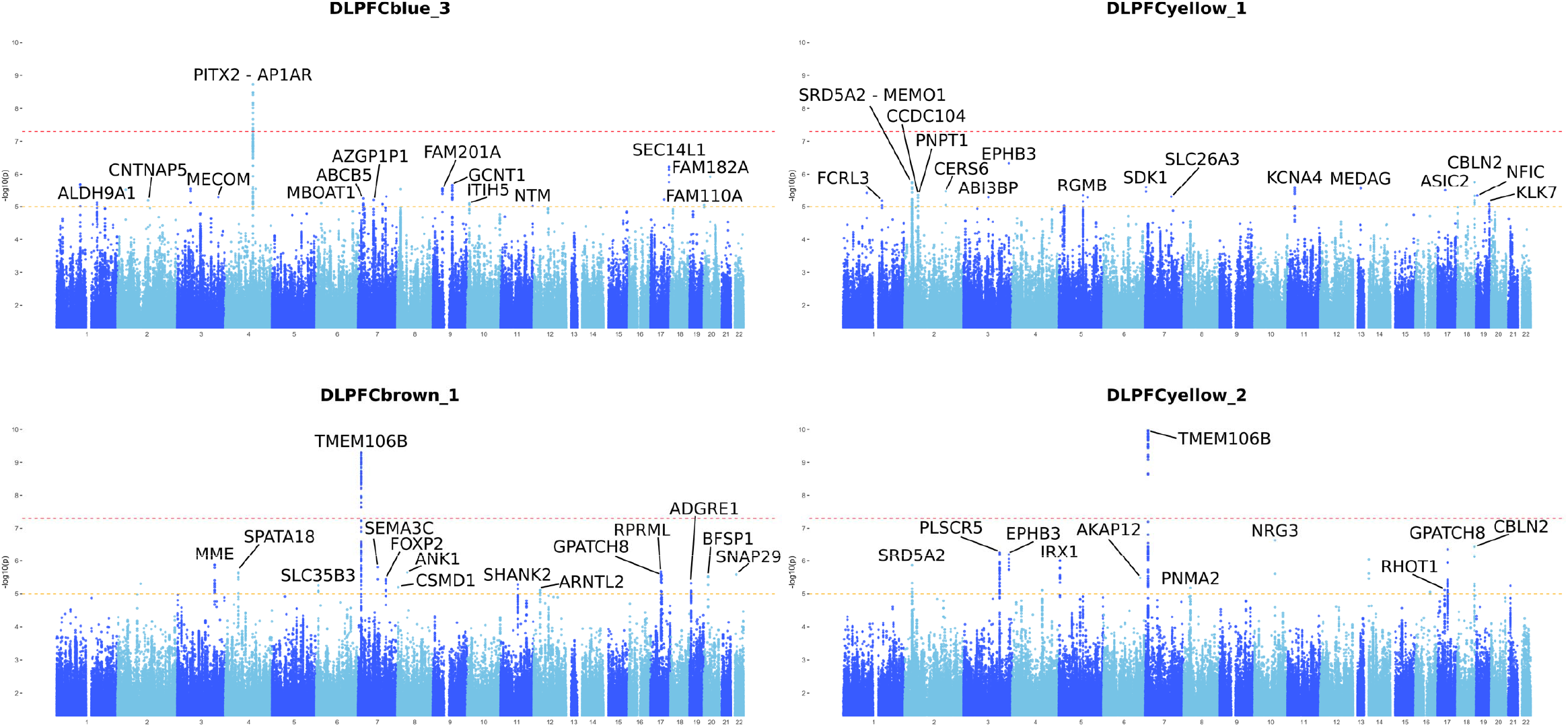
Manhattan plots of single-variant association of select submodule eigengenes in ROSMAP. Eigengene expression for each submodule was used as a quantitative trait when performing single-variant mapping. These Manhattan plots were generated for select DLPFC region submodule eigengenes. Multiple submodule eigengenes were associated with SNPs at a genome-wide significance level of p = 5e-08 (red dotted line). Loci of interest are annotated with the gene closest to the region. Some SNPs were also detected at a genome-wide suggestive level of p = 1e-05 (yellow dotted line). DLPFCblue_3 contains genes related to the TREM2/TYROBP pathway, an important network of genes related to microglial activation during neuroinflammation of the brain. Submodules were associated with both unique and overlapping loci. For example, DLPFCbrown_1 and DLPFCyellow_2 are derived from separate co-expression modules but were both associated with the *TMEM106B* locus. Similarly, DLPFCyellow_1 and DLPFCyellow_2 were derived from the same co-expression module but were associated with a mix of overlapping and unique loci.

**Figure 4:**
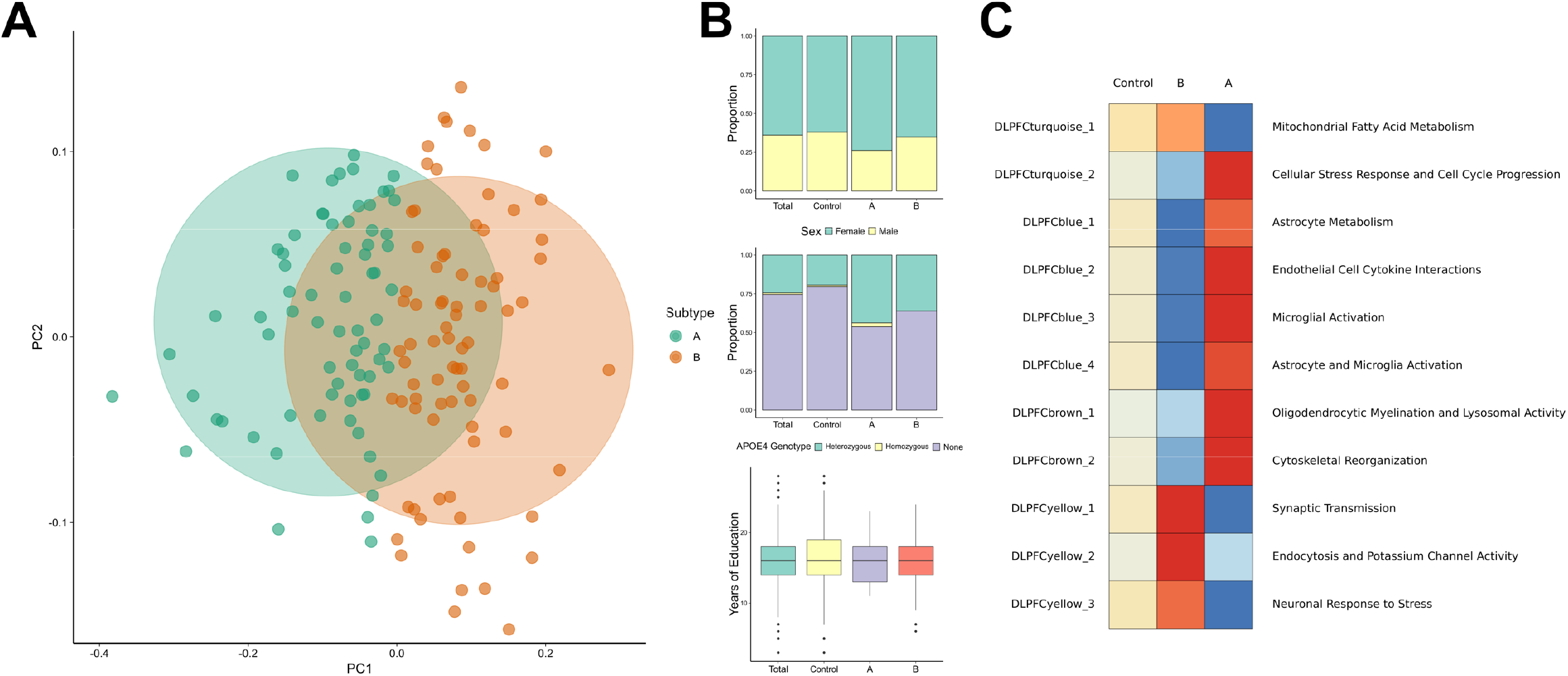
Clustering on eigengene expression in ROSMAP data generates 2 subtypes. (A) Eigengene expression was used to cluster cases into subtypes using K-Means clustering for the DLPFC region. The number of clusters were determined by democratizing results across 30 mathematical indices using the NbClust R package. Two clusters with relatively equal number of cases were generated. (B) There were no significant differences in proportion of sex, *APOEε4* genotype, and years of education between subtypes. (C) The scaled eigengene expression profile of the subtypes demonstrated a strong immune and neuronal signal when compared to control and MCI decedents.

Eigengene expression profiles for each subtype were used to assess the association of each subtype with molecular and biological pathways associated with submodules. An example for the ROSMAP cohort is shown in Figure 4. We observed no significant enrichment of cognitive or neuropathological measures between the subtypes for the DLPFC region (Figure S6).

### ROSMAP subtypes differ in inflammatory response

In order to better understand the underlying molecular differences across the novel LOAD associated subtypes in the ROSMAP cohort and to identify potential subtype specific candidate markers, differential expression analysis was performed for each of the previously defined subtypes against a set of controls (Figure 5a). Each of the two subtypes was compared to a set of 471 decedents from the ROSMAP cohort that were either cognitively normal or had mild cognitive impairment. The Venn diagram in Figure 5b depicts the comparison across the different subtypes. Interestingly, cases associated with Subtype A showed a stronger transcriptional response with 127 differentially expressed genes (adjusted p-values < 0.05, absolute log fold change > 0.5) when compared with controls. Of these genes, 86 were up-regulated and 41 were down-regulated. Among the most significantly down-regulated genes associated with Subtype A cases was the stress-response mediator corticotropin-releasing hormone *(CRH).* Overacting *CRH* signaling has been implicated in inflammatory disorders and LOAD where it has been proposed as a therapeutic target to reduce the negative effects of chronic stress related to memory function and amyloid beta (Aβ) production^30^. Cases associated with Subtype B had 40 differentially expressed genes (adjusted p-values < 0.05, absolute log fold change > 0.5), 39 of which were down-regulated when compared to controls. Notably, two key pro-inflammatory mediators of amyloid deposition *(S100A8, S100A9)* were among the most significantly down-regulated genes in Subtype B decedents when compared to controls (Figure 5a). Both genes, which are established inflammatory biomarkers, are part of a complex that serves as a critical link between the amyloid cascade and inflammatory events in LOAD^31^. Furthermore, multiple pathways linked to *S100A8/9* activation, including IL-10 signaling and complement activation were enriched across down-regulated genes in Subtype B but not in Subtype A decedents as highlighted in Figure 5c. In addition, molecular pathways linked to microglia activation (Figure S8), the immune response, and the stress response were found among the most significant pathways and gene sets (Table S13, Table S14) that differ across subtypes. Gene set enrichment analysis revealed a subset of genes linked to the KEGG osteoclast differentiation pathway (Figure S8), including known AD risk markers such as *TREM2, TYROBP*, and *CCL2* among others which were highly up-regulated in Subtype A cases compared to Subtype B cases. This highlights that both molecularly defined LOAD subtypes differ in their immune response and that known LOAD biomarkers, including *S100A8/A9^32^, TREM2*, and *CCL2* might be used to stratify patients based upon their inflammatory response to the observed disease state. These results were consistent with the functional annotations of the previously defined submodules that define both subtypes (Figure 4C).

**Figure 5:**
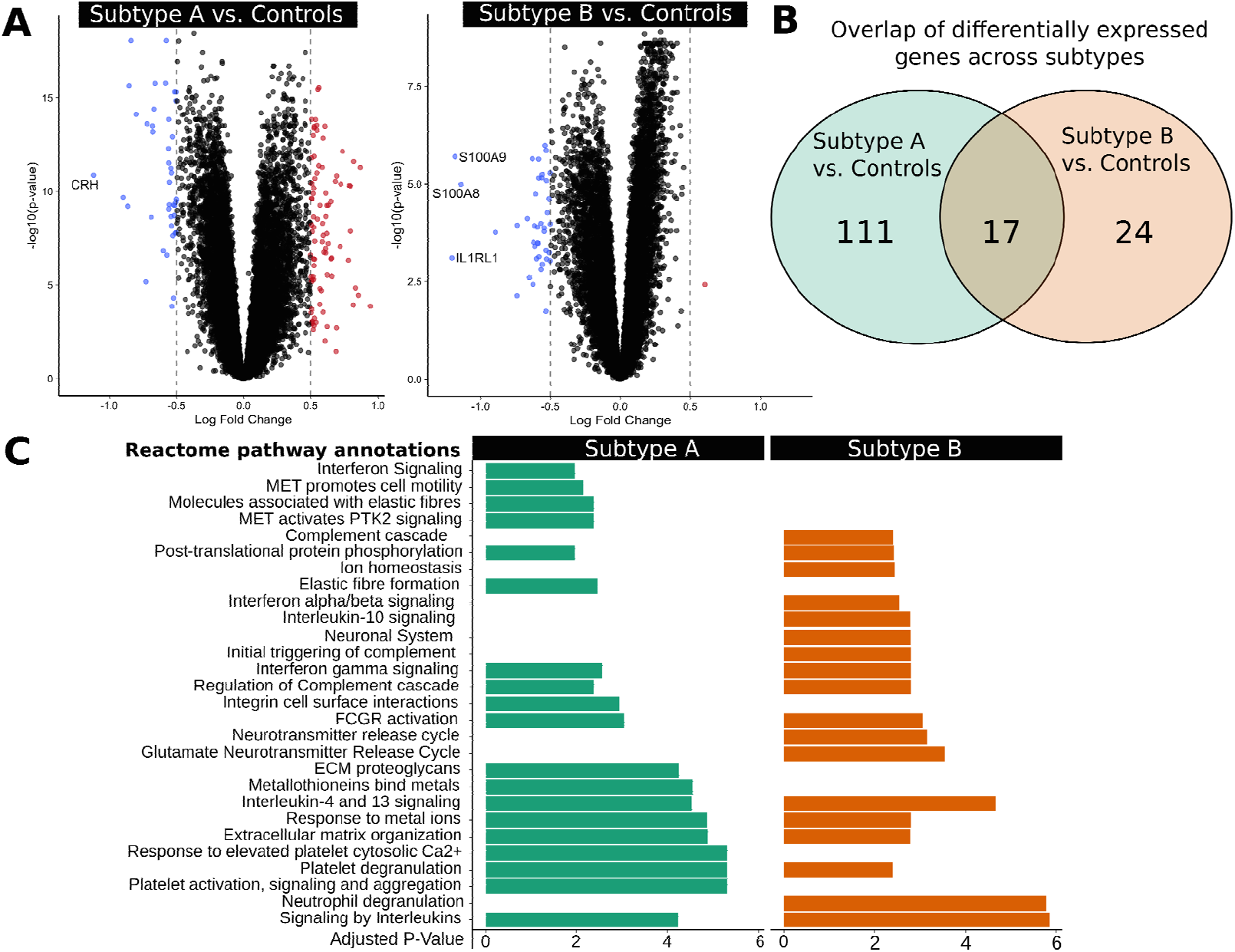
Differential expression analysis of ROSMAP subtypes reveals heterogeneity in inflammatory response in LOAD cases. (A) Differential expression analysis comparing each subtype to control decedents for the DLPFC region was performed using the limma R package. We show up-regulated (red, p < 0.05, log fold change >0.5) and down-regulated (blue, p < 0.05, log fold change < −0.5) genes in the volcano plot and label genes that have an absolute log fold change of greater than 1 (dotted lines). (B) Differentially expressed genes (p < 0.05, absolute log fold change > 0.5) from the analysis show a partial overlap between subtypes. (C) Top Reactome pathways for differentially expressed genes for both subtypes are reported. Subtype A demonstrates an enrichment of immune and stress-response related pathways across up-regulated genes, while Subtype B demonstrates a down-regulation of a set of specific immune-related pathways linked to S100A8/A9 activation.

### Single variant association mapping for ROSMAP decedents

Genome wide association mapping revealed a differential enrichment of significant variants across subtypes (Figure 6, Table S9, Table S10, Table S11, Table S12). Loci were associated with one or more submodule eigengenes, as shown in Figure 6. One genome-wide suggestive allele in *TMEM106B* was identified for Subtype B (p-value < 4×10^−6^, rs1990620^G^). This association was replicated at a genome-wide suggestive level in association with the DLPFCbrown_2 eigengene and at a genome-wide significant level with the DLPFCbrown_1 and DLPFCyellow_2 eigengenes (Figure 3). DLPFCbrown_1 contains genes related to myelination and lysosomal activity (KEGG pathways hsa00600 and hsa04142), while DLPFCyellow_2 contains genes related to endocytosis and potassium channel activity (KEGG pathway hsa04144 and Reactome pathway R-HSA-1296071). *TMEM106B* is a known modifier of neurodegenerative disease and cognitive aging, which has been previously linked with cognitive performance^33^. Loss of *TMEM106B* function has been shown to rescue lysosomal phenotypes related to frontotemporal dementia^34^. The identified protective allele rs1990620^G^ is a known CCCTC-binding factor (CTCF) site, which has been shown to modify the inflammatory response in the course of aging^35^. Besides the association with *TMEM106B* in Subtype B, protective variants near *MTUS2* were identified which are in close vicinity to *HMGB1*, a locus that has been previously implicated in brain atrophy^36^. A differential expression analysis of haplotype carriers of the protective rs1990620^G^ variant in *TMEM106B* showed an up-regulation of neuroactive ligand receptor interactions, while decedents carrying the risk variant showed significant up-regulation for pathways related to Osteoclast differentiation (KEGG pathway hsa04380) and neuroinflammation (data not shown).

**Figure 6:**
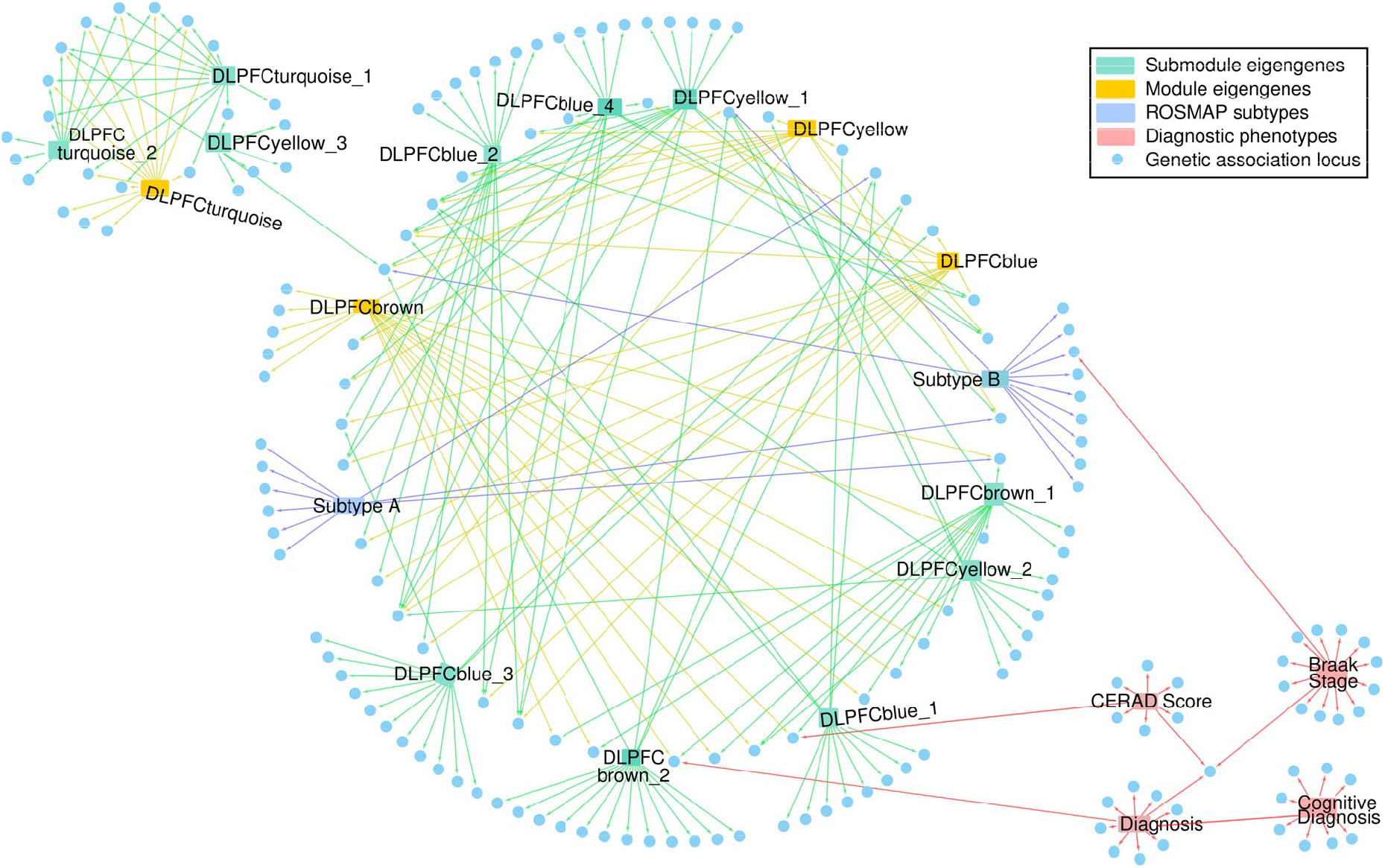
Network of phenotypes and associated loci. We created a directed network describing the loci detected from the multiple analyses in this study. Blue nodes represent loci associated with a phenotype. Red nodes represent phenotypes. An edge from a phenotype to a genetic locus signifies that the locus is associated with the specified phenotype. Diagnostic phenotypes (red edges) were associated with some of the loci detected in this study. The module eigengenes (yellow edges), submodule eigengenes (green edges), and subtypes (blue edges) were associated with overlapping and unique loci (center and left). A community of loci was associated with multiple submodules associated with microglia, endothelial cells, astrocytes, and oligodendrocytes (center). A small community of loci was associated with submodules related to proteostasis (left). Diagnostic phenotypes included CERAD scores, Braak stages, cognitive diagnosis, and case-control association.

### Suggestive SNPs in ROSMAP are replicated in other cohorts

A total of 1326 unique SNPs representing 163 loci were genome-wide suggestive or significant (p-value < 1×10^−5^) in the DLPFC region when pooled from all 11 DLPFC eigengenes and two subtype-specific variant mapping analyses. Of these, 645 SNPs were replicated in the PHG analyses, 762 SNPs were replicated in the FP analysis, and 482 SNPs were replicated in the TCX analyses (p-value < 1×10^−2^). The *TMEM106B* variant associated with dementia, rs1990620, was replicated in all cohorts. Of the 163 loci, 29 loci across 27 studies had been previously reported in the NHGRI-EBI catalog such that the most significant SNP from the prior study was a suggestive SNP in the DLPFC region (Table S15, Table S16, Table S17, Table S18).

## DISCUSSION

Common complex diseases such as LOAD are characterized by phenotypic heterogeneity and the presence of multiple common variants affecting disease risk. In this study, we present an analysis that uses transcriptomic co-expression data and whole-genome sequencing from multiple cohorts to dissect phenotypic heterogeneity and identify potential genetic drivers of complex trait pathology in LOAD.

Here, we used an iterative pruning approach based on 26 human post-mortem co-expression modules to generate 68 novel submodules that contained genes associated with LOAD specific biological pathways and molecular processes. Indeed, we observed that genes in the novel submodules are enriched for functional terms that were specific to pathways associated with LOAD, such as lipid modification, the TREM2/TYROBP pathway, and tau-protein kinase activity. Furthermore, submodules from all six brain regions clustered independently of the co-expression module of origin and brain region, suggesting that the genes captured in each submodule represented signals that were associated with LOAD pathology rather than cohort- or tissue-specific factors. Notably, submodules were much more specific for markers of different brain cell types, suggesting that the processes associated with submodules represent the pathological signals from these specific cell types. This is in line with recent studies showing that different cell types in the brain play specific roles at different stages in the pathogenesis of LOAD^37^. Taken together, our results demonstrate that the novel human co-expression submodules identified in this study capture cell-type specific pathways associated with LOAD pathogenesis in the brain.

Mapping the eigengene expression for individual submodules represents a pathway- or process-level alternative to expression quantitative trait locus (eQTL) mapping for each individual transcript. Since the human co-expression submodules represented pathological, cell-type specific pathways in LOAD brain tissue, mapping eigengene expression for decedents was expected to identify genetic drivers of LOAD pathology. RNA-Seq data from post-mortem brain tissue in human cohorts contains a strong immune signal, as evidenced by repeated identification of genetic loci related to microglial response in meta-analyses with increasingly large cohorts^5,38^. Using submodule eigengenes as quantitative traits for single-variant association provided an opportunity to identify genetic drivers of biological processes that are known to be drivers of early LOAD pathogenesis, such as astrogliosis, neuronal plasticity, myelination, and vascular blood brain barrier interactions^37^. Suggestive variants identified were unique to subsets of submodules. For instance, the *TMEM106B* locus was associated at a genome-wide significant level with the DLPFCbrown_1 and DLPFCyellow_2 eigengenes (Figure 3), representing processes related to oligodendrocytic myelination, lysosomal activity, endocytosis, and potassium channel activity. The *TMEM106B* locus has been implicated in cognitive aging, with functional consequences in frontotemporal dementia related to lysosomal activity^33–55^. A submodule of particular interest is the microglia-associated submodule DLPFCblue_3, which contains genes related to the TREM2/TYROBP cascade. The *FAM110A* locus is close to rs1014897 and the *CNTNAP5* locus is close to rs76854344, both variants have been previously associated with posterior cortical atrophy and LOAD^39^. The *NTM* locus is close to rs1040103, a variant that has been associated with white blood cell count^40^. Thus, quantitative trait mapping of single variants using eigengene expression for submodules presented in this study can elucidate genetic factors specific to associated pathological pathways.

Furthermore, eigengenes represent a dimensional reduction of transcriptomic data onto axes of pathological relevance. Thus, we expected that clustering on the eigengene expression of LOAD cases would generate pathway-level profiles of putative molecular LOAD subtypes based on case heterogeneity. As anticipated, we observed that average eigengene expression was enriched by subtype for multiple submodules in all four brain regions tested. Strikingly, these enrichments were diametric in the subtypes generated for LOAD cases, an example of which is presented for the DLPFC region in Figure 4. Similar enrichment patterns were identified in the other three brain regions. These results suggest that the biological programs identified by submodules in this study align themselves along the heterogeneity of transcriptomic data present in LOAD cases across multiple cohorts rather than differentiating solely based on cases and controls. Furthermore, the stratification of patients based on submodule expression profiles demonstrated that there is significant variation in immune response in post-mortem brain tissue, a process that is considered a hallmark of LOAD pathogenesis (Figure 5, Figure S8). Variants associated with the subtype specificity metric overlapped with the variants associated with individual submodule eigengenes (Figure 6). This suggests that the genetic factors that influenced subtypes can be dissected into loci driving specific submodules. Furthermore, the deconstruction of genetic loci can provide the basis for more targeted treatment of dysfunctional pathways that contribute to different subtypes of LOAD.

Our subtypes in the DLPFC brain region of the ROSMAP cohort represent differences in transcriptomic profiles of LOAD cases derived from post-mortem RNA-Seq data. A lack of temporal data makes it challenging to decisively interpret these profiles. The subtypes may represent distinct LOAD endpoints, differences in disease severity, environmental effects, or phases of molecular pathology. Neither subtype was associated with cognitive or neuropathological outcome (Figure S6). Furthermore, covariates such as sex, *APOE* genotype, and years of education were not significantly enriched in any given subtype (Figure 4). This suggests that the transcriptomic profiles do not represent transitions in disease severity and that there are overall risk factors not reflected in transcriptomic subtypes. Furthermore, both subtypes are associated with unique loci that belong to the same community of loci detected by submodule mapping (Figure 6), indicating that the subtypes capture various combinations of genetic elements that lead to LOAD pathology. While suggestive, these transcriptomic LOAD subtypes will require further validation in cohorts that adequately control for disease progression.

The methodology presented in this study is not limited to RNA-Seq data and can be performed on other omics, such as proteomics or metabolomics. As such data become available for the decedents in these cohorts, this analysis can be expanded across these additional informative dimensions.

## ACKNOWLEDGEMENTS

This study was supported by the National Institutes of Health grant U54 AG 054354. Additional funding was provided by the Barbara H. Sanford Endowed Scholarship Fund, the Robert E. Garrity Cooperative Education Fund as well as the Park and Goldwater Scholarship funds. We thank A. Saykin and K. Nho for helpful conversations.

The results published here are in whole or in part based on data obtained from the AMP-AD Knowledge Portal (<u>doi:10.7303/syn2580853</u>). Study data were provided by the Rush Alzheimer’s Disease Center, Rush University Medical Center, Chicago. Data collection was supported through funding by NIA grants P30AG10161, R01AG15819, R01AG17917, R01AG30146, R01AG36836, U01AG32984, U01AG46152, the Illinois Department of Public Health, and the Translational Genomics Research Institute.

The results published here are in whole or in part based on data obtained from the AMP-AD Knowledge Portal (<u>doi:10.7303/syn2580853</u>). Study data were provided by the following sources: The Mayo Clinic Alzheimer’s Disease Genetic Studies, led by Dr. Nilufer Taner and Dr. Steven G. Younkin, Mayo Clinic, Jacksonville, FL using samples from the Mayo Clinic Study of Aging, the Mayo Clinic Alzheimer’s Disease Research Center, and the Mayo Clinic Brain Bank. Data collection was supported through funding by NIA grants P50 AG016574, R01 AG032990, U01 AG046139, R01 AG018023, U01 AG006576, U01 AG006786, R01 AG025711, R01 AG017216, R01 AG003949, NINDS grant R01 NS080820, CurePSP Foundation, and support from Mayo Foundation. Study data includes samples collected through the Sun Health Research Institute Brain and Body Donation Program of Sun City, Arizona. The Brain and Body Donation Program is supported by the National Institute of Neurological Disorders and Stroke (U24 NS072026 National Brain and Tissue Resource for Parkinson’s Disease and Related Disorders), the National Institute on Aging (P30 AG19610 Arizona Alzheimer’s Disease Core Center), the Arizona Department of Health Services (contract 211002, Arizona Alzheimer’s Research Center), the Arizona Biomedical Research Commission (contracts 4001, 0011, 05-901 and 1001 to the Arizona Parkinson’s Disease Consortium) and the Michael J. Fox Foundation for Parkinson’s Research. The results published here are in whole or in part based on data obtained from the AMP-AD Knowledge Portal (<u>doi:10.7303/syn2580853</u>). These data were generated from postmortem brain tissue collected through the Mount Sinai VA Medical Center Brain Bank and were provided by Dr. Eric Schadt from Mount Sinai School of Medicine.

## AUTHOR’S CONTRIBUTIONS

NM, CP, AH, GA and CJ performed the genetic and transcriptomic analysis. SS annotated the functional variants. SM and PC provided cognitive and phenotype data for the analysis. BL and AT provided additional transcriptomic data for the analysis. GWC supervised and designed the project. NM, CP and GWC wrote the manuscript. All authors read and approved the final manuscript.

## CONSENT FOR PUBLICATION

All authors have approved of the manuscript and agree with its submission.

