## Supplemental Information for "Transcriptomic Stratification of Late-Onset Alzheimer’s Cases Reveals Novel Genetic Modifiers of Disease Pathology"

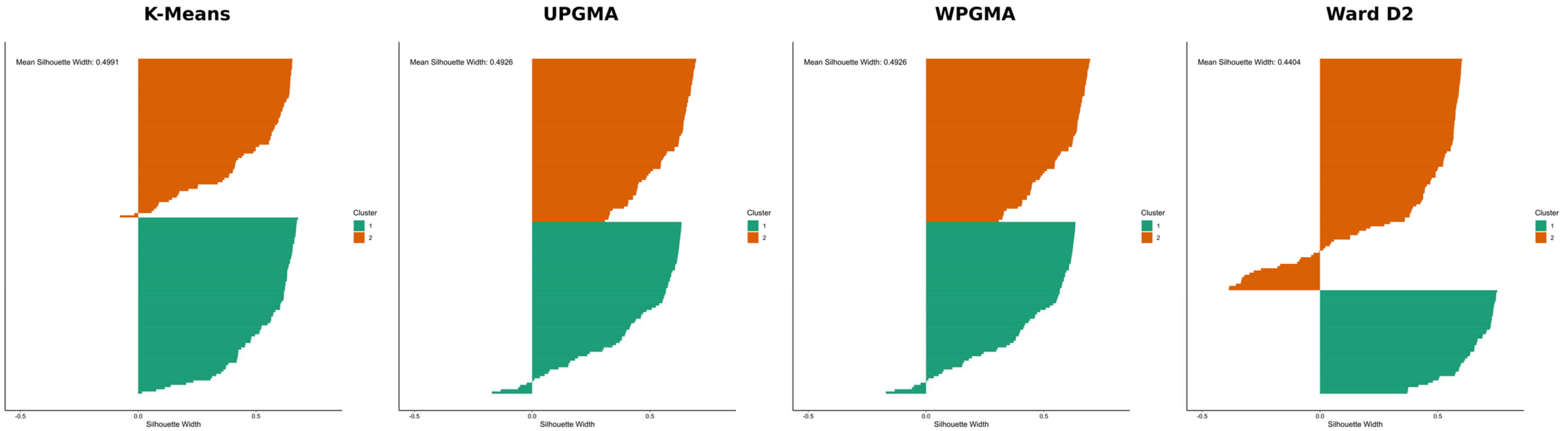


**Supplemental Figure 1: Silhouette plots generated by four different clustering methods used on LOAD cases from the ROSMAP cohort.** A silhouette is a measure that simultaneously characterizes the similarity of an object to its cluster and the dissimilarity of the object with other clusters. The mean silhouette width of a cluster represents how similar objects are to the centroid of the cluster, and the mean silhouette width of all objects represent how well the data have been clustered. In the case of ROSMAP, the K-means method had the highest mean silhouette width across all LOAD cases. We performed a similar analysis for other stratified brain regions.

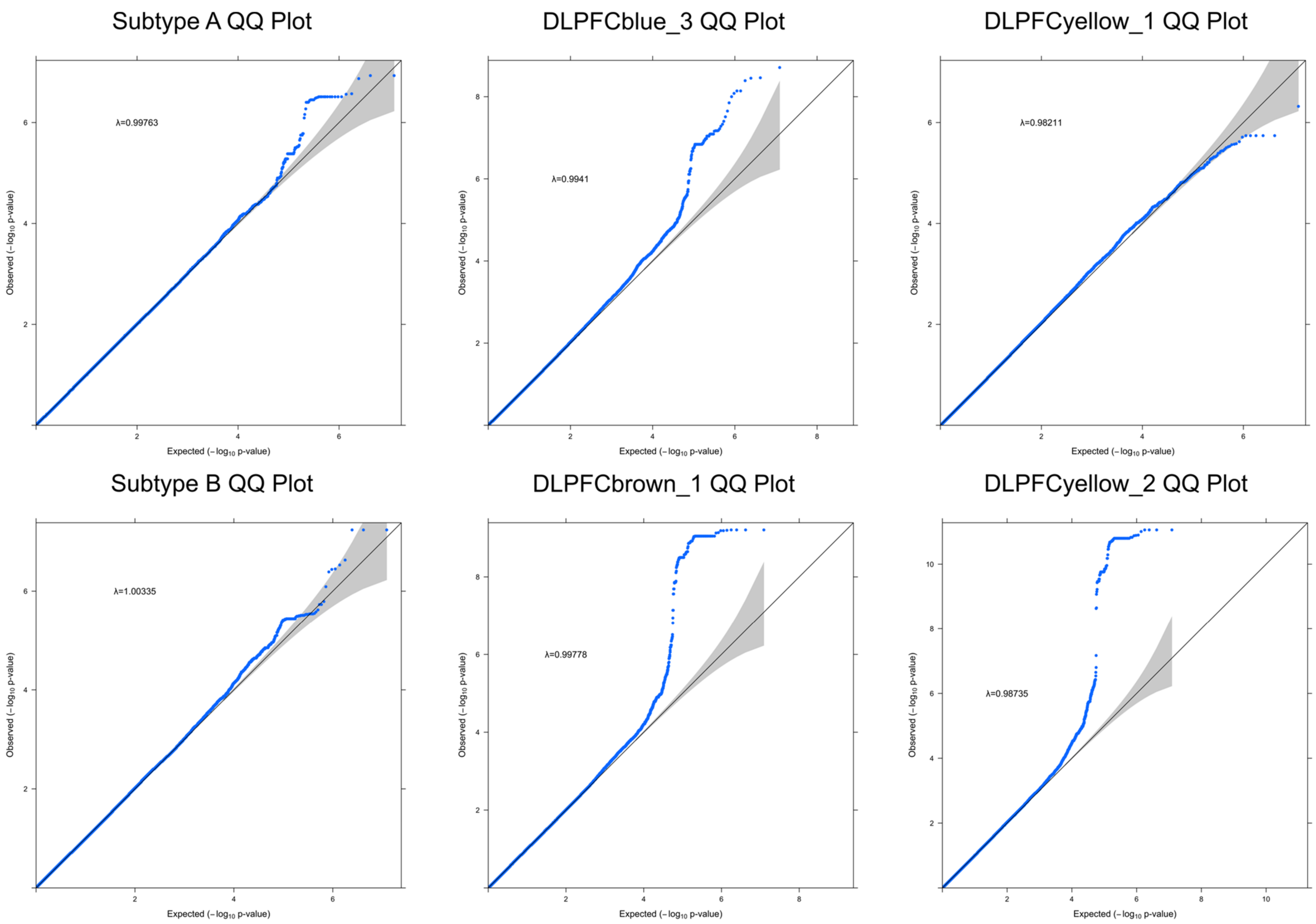


**Supplemental Figure 2: QQ Plots of observed p-values from single-variant association in the ROSMAP cohort.** QQ plots of select single-variant association analyses of the DLPFC region that were presented in Figure 3 and Figure S6 show that there is minimal genomic inflation, and consequently, minimal population substructure effects on the analyses. The genomic inflation factor for each QQ plot is also reported. Each QQ plot compares the expected and observed distribution of p-values obtained from the association analysis for a given phenotype.

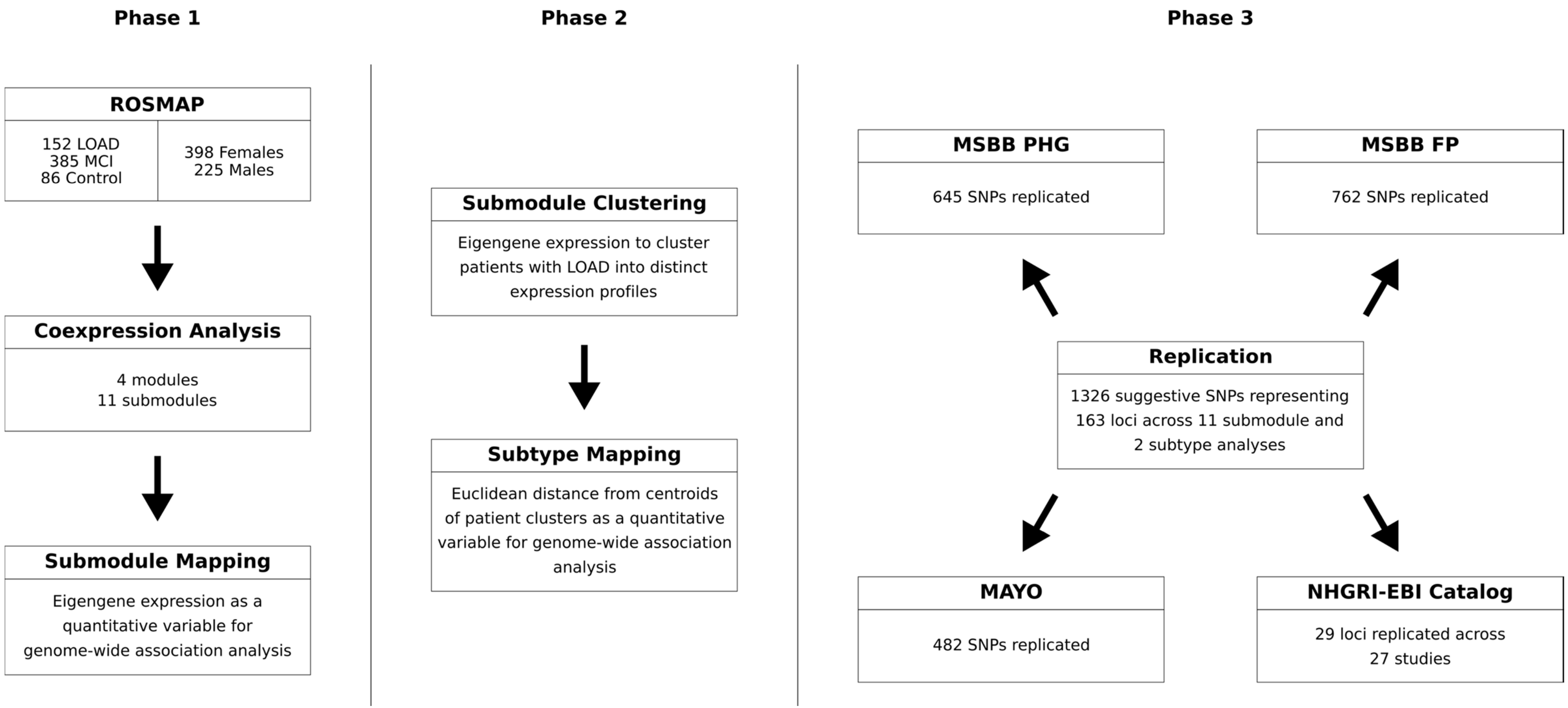


**Supplemental Figure 3: The complete analysis carried out in this study is divided into three phases**. Phase 1 involved the co-expression analysis of ROSMAP and other cohorts to generate submodules representing biological processes involved in Alzheimer's pathology. Eigengene expression from the submodules were used to perform single-variant association and identify loci that act as putative genetic drivers of these biological pathways. Phase 2 involved the clustering of LOAD cases in ROSMAP and other cohorts based on an agnostic clustering method. Subtypes were mapped using single-variant association to identify loci that may explain the heterogeneity observed in LOAD cases. Phase 3 involved the replication of genome-wide suggestive or genome-wide significant SNPs from the ROSMAP cohort in other tissue regions and previous studies. SNPs were replicated in three other tissue regions (PHG, FP, TCX) and in 27 studies from the NHGRI-EBI catalog.

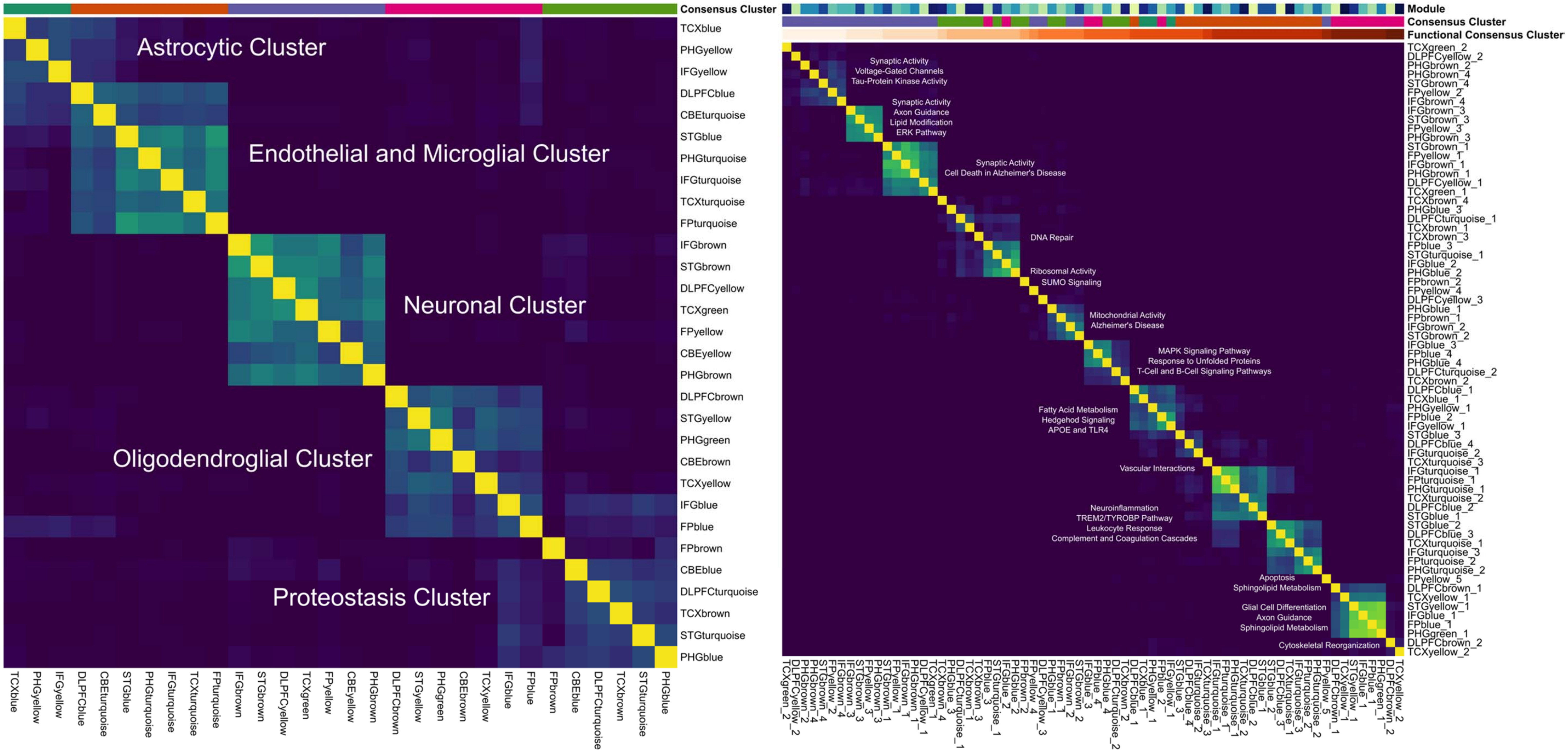


**Supplemental Figure 4: Clusters of modules and submodules based on gene overlap reveal cell-type and functional signatures.** (A) A previous study by Logsdon *et al.* reported 5 consensus clusters across 7 tissue regions based on the modules generated for each tissue region. A Jaccard matrix heatmap is used to visualize the overlap of genes in each module between tissue regions and cohorts. (B) Submodules were divided into 15 functional clusters based on hierarchical clustering that demonstrated specificity for certain biological pathways. These functional clusters formed independently of module of origin and tissue of origin.

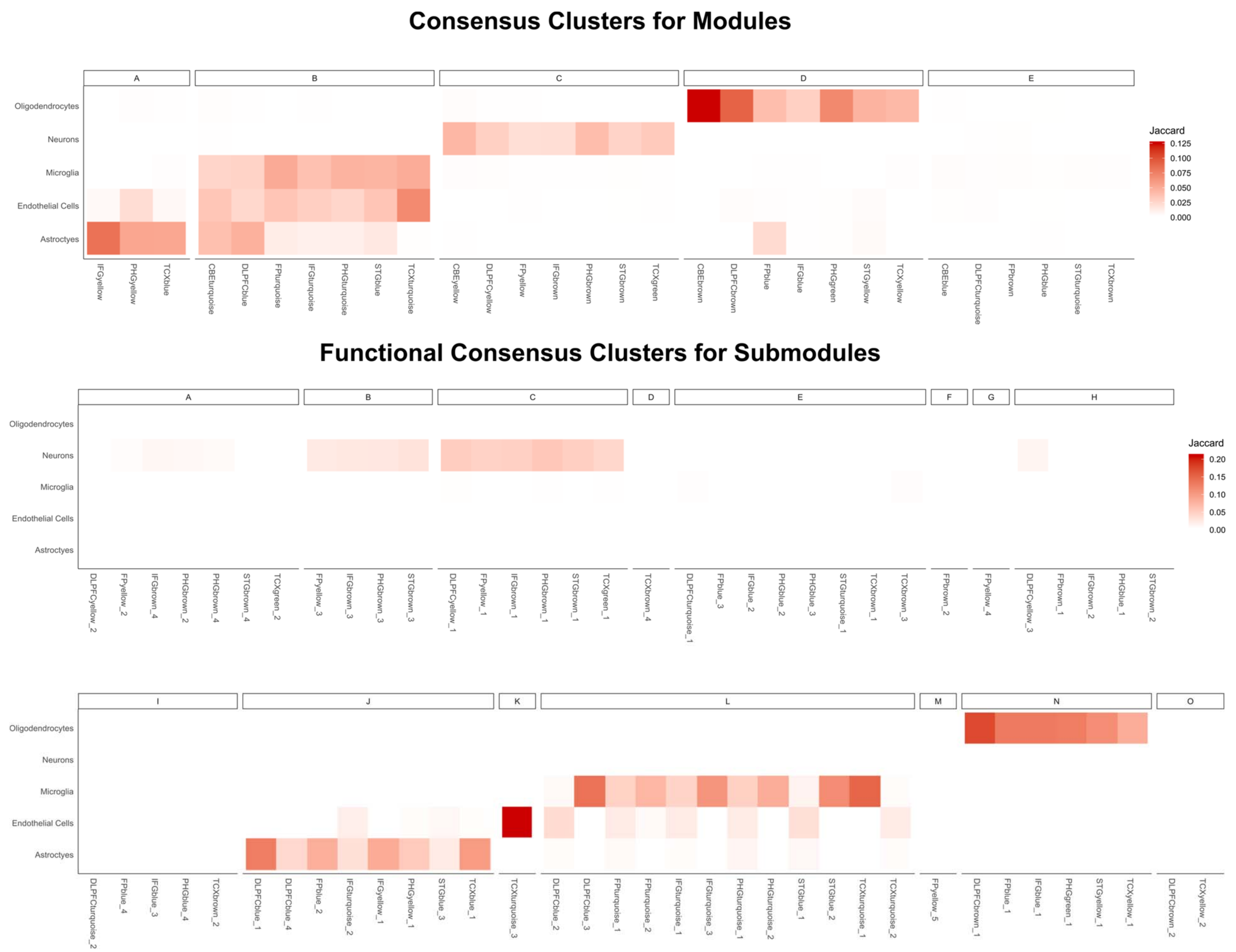


**Supplemental Figure 5: Functional consensus clusters demonstrate cell-type specificity.** Brain tissue cell-type specific markers reported previously by McKenzie *et al.* were used to assess the cell-type specificity of modules and submodules. Consensus clusters B broadly captured astrocytic, endothelial, and microglial signals. This signal was resolved in the functional consensus clusters generated using the submodules across functional consensus clusters J, K, and L.

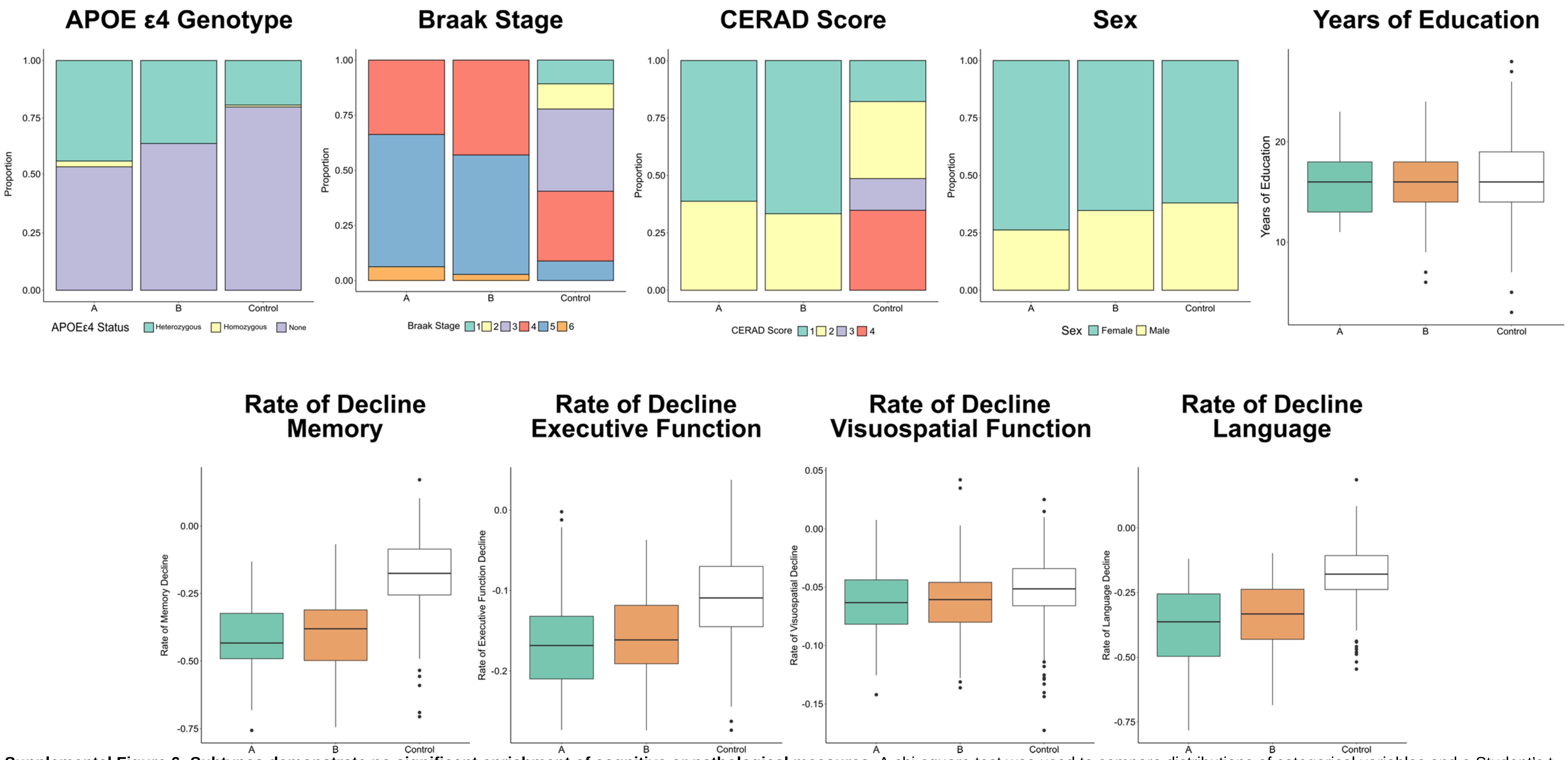


**Supplemental Figure 6: Subtypes demonstrate no significant enrichment of cognitive or pathological measures.** A chi-square test was used to compare distributions of categorical variables and a Student's t-test was used to compare distributions of quantitative variables between subtypes ($\alpha = 0.05$ significance level). Braak stages are a measure of neurofibrillary tangles and CERAD scores are a measure of neuritic plaques. Rates of decline in cognitive phenotypes were measured previously by Mukherjee *et al.* for a subset of the ROSMAP cohort.

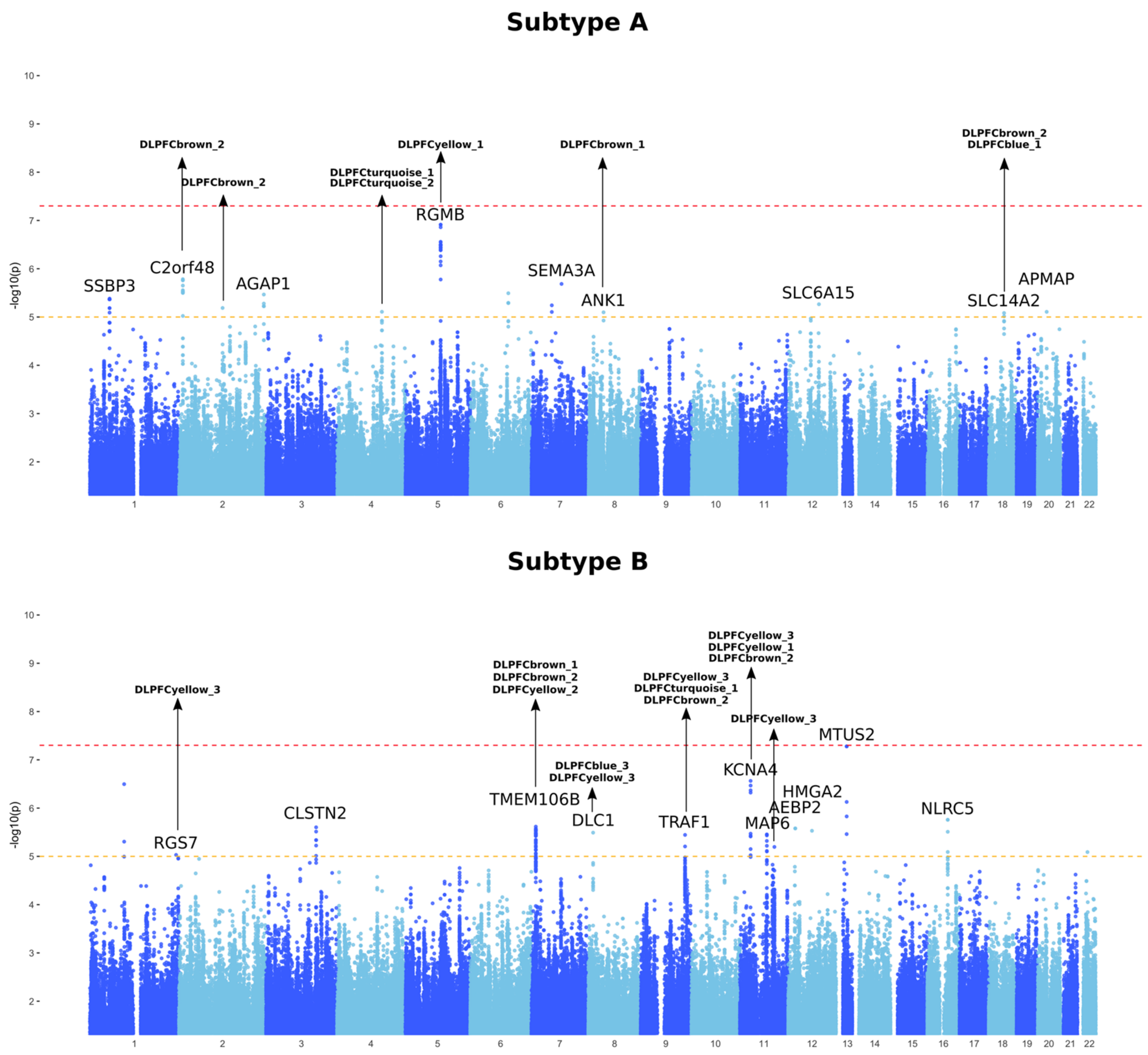


**Supplemental Figure 7: Manhattan plots of single-variant association of the subtype specificity metric in ROSMAP**
Single-variant association of the subtype specificity metric of the two subtypes in the DLPFC region recapitulate multiple loci generally detected at a higher power with submodule eigengenes. Certain loci, such as *MTUS2*, were not detected in previous submodule eigengene associations.

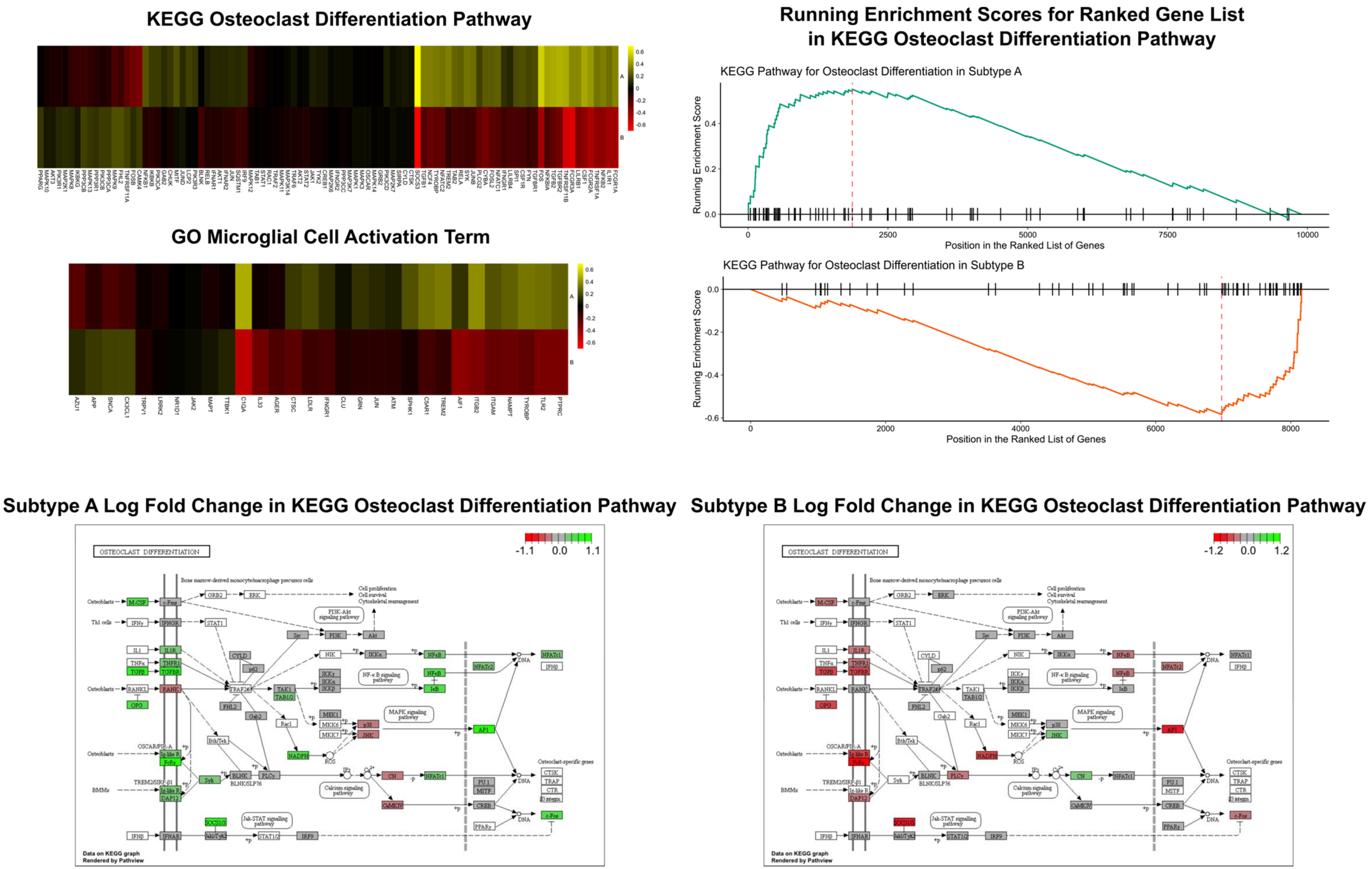


**Supplemental Figure 8: Pathway enrichment analysis of differentially expressed genes in subtypes from the ROSMAP cohort.** Pathway enrichment analyses of subtypes generated using the DLPFC region data show upregulation of the TREM2/TYROBP pathway in Subtype A and downregulation of the pathway in Subtype B. The KEGG Osteoclast Differentiation pathway and GO Microglial Cell Activation term contain many of the genes associated with the TREM2/TYROBP pathway.

| Cohort | Tissue | Decedents |
|---|---|---|
| Mayo | Temporal Cortex | 262 |
| MSBB | Frontopolar Prefrontal Cortex | 214 |
| MSBB | Inferior Temporal Gyrus | 187 |
| MSBB | Parahippocampal Gyrus | 160 |
| MSBB | Superior Temporal Gyrus | 187 |
| ROSMAP | Dorsolateral Prefrontal Cortex | 623 |

**Supplemental Table 1: Summary of Cohorts**

RNA-Seq and whole genome sequencing data from the Mayo Clinic, the Mount Sinai Brain Bank, and the Rush University's Religious Orders Study and Memory and Aging Project. 6 brain regions from these studies were used. The number of RNA-Seq samples and whole genome sequencing data for each tissue are reported.

| Cohort | Brain Region | Diagnosis | Sex | Decedents |
|---|---|---|---|---|
| Mayo | TCX | TCX.AD | F | 49 |
| Mayo | TCX | TCX.AD | M | 31 |
| Mayo | TCX | TCX.OTHER | F | 50 |
| Mayo | TCX | TCX.OTHER | M | 61 |
| Mayo | TCX | TCX.CONTROL | F | 35 |
| Mayo | TCX | TCX.CONTROL | M | 36 |
| MSBB | FP | FP.AD | F | 63 |
| MSBB | FP | FP.AD | M | 27 |
| MSBB | FP | FP.OTHER | F | 54 |
| MSBB | FP | FP.OTHER | M | 25 |
| MSBB | FP | FP.CONTROL | F | 23 |
| MSBB | FP | FP.CONTROL | M | 22 |
| MSBB | IFG | IFG.AD | F | 55 |
| MSBB | IFG | IFG.AD | M | 24 |
| MSBB | IFG | IFG.OTHER | F | 50 |
| MSBB | IFG | IFG.OTHER | M | 21 |
| MSBB | IFG | IFG.CONTROL | F | 17 |
| MSBB | IFG | IFG.CONTROL | M | 20 |
| MSBB | PHG | PHG.AD | F | 46 |
| MSBB | PHG | PHG.AD | M | 18 |
| MSBB | PHG | PHG.OTHER | F | 39 |
| MSBB | PHG | PHG.OTHER | M | 19 |
| MSBB | PHG | PHG.CONTROL | F | 18 |
| MSBB | PHG | PHG.CONTROL | M | 20 |
| MSBB | STG | STG.AD | F | 54 |
| MSBB | STG | STG.AD | M | 27 |
| MSBB | STG | STG.OTHER | F | 49 |
| MSBB | STG | STG.OTHER | M | 20 |
| MSBB | STG | STG.CONTROL | F | 20 |
| MSBB | STG | STG.CONTROL | M | 17 |
| ROSMAP | DLPFC | DLPFC.AD | F | 106 |
| ROSMAP | DLPFC | DLPFC.AD | M | 46 |
| ROSMAP | DLPFC | DLPFC.OTHER | F | 245 |
| ROSMAP | DLPFC | DLPFC.OTHER | M | 140 |
| ROSMAP | DLPFC | DLPFC.CONTROL | F | 47 |
| ROSMAP | DLPFC | DLPFC.CONTROL | M | 39 |

**Supplemental Table 2: Summary of Cohorts by Diagnosis and Sex**

For each of the six brain regions, possible diagnoses include Late-Onset Alzheimer's Disease (AD), unaffected elderly controls (CONTROL), and other decedents (OTHER). In MSBB and ROSMAP, other decedents were diagnosed with mild cognitive impairment while other decedents in Mayo were diagnosed with either progressive supranuclear palsy (PSP) or pathological aging (PA).

| Cohort | Brain Region | Module | Genes |
|---|---|---|---|
| Mayo | TCX | TCXgreen | 2766 |
| Mayo | TCX | TCXyellow | 2013 |
| Mayo | TCX | TCXbrown | 1851 |
| Mayo | TCX | TCXblue | 1713 |
| Mayo | TCX | TCXturquoise | 1131 |
| MSBB | FP | FPyellow | 4426 |
| MSBB | FP | FPblue | 1991 |
| MSBB | FP | FPbrown | 1289 |
| MSBB | FP | FPturquoise | 1001 |
| MSBB | IFG | IFGbrown | 4673 |
| MSBB | IFG | IFGblue | 2885 |
| MSBB | IFG | IFGturquoise | 1456 |
| MSBB | IFG | IFGyellow | 743 |
| MSBB | PHG | PHGblue | 3733 |
| MSBB | PHG | PHGbrown | 2123 |
| MSBB | PHG | PHGturquoise | 1195 |
| MSBB | PHG | PHGgreen | 1151 |
| MSBB | PHG | PHGyellow | 910 |
| MSBB | STG | STGbrown | 3414 |
| MSBB | STG | STGturquoise | 2404 |
| MSBB | STG | STGyellow | 1799 |
| MSBB | STG | STGblue | 1171 |
| ROSMAP | DLPFC | DLPFCyellow | 3019 |
| ROSMAP | DLPFC | DLPFCturquoise | 2489 |
| ROSMAP | DLPFC | DLPFCblue | 1751 |
| ROSMAP | DLPFC | DLPFCbrown | 882 |

**Supplemental Table 3: Summary of Modules**

Modules were generated independently for each tissue region. The number of genes in each module are reported. 26 modules were used in this study.

| Cohort | Brain Region | Module | Submodule | Genes |
|---|---|---|---|---|
| Mayo | TCX | TCXblue | TCXblue_1 | 715 |
| Mayo | TCX | TCXbrown | TCXbrown_1 | 391 |
| Mayo | TCX | TCXbrown | TCXbrown_2 | 251 |
| Mayo | TCX | TCXbrown | TCXbrown_3 | 245 |
| Mayo | TCX | TCXbrown | TCXbrown_4 | 120 |
| Mayo | TCX | TCXgreen | TCXgreen_1 | 2090 |
| Mayo | TCX | TCXgreen | TCXgreen_2 | 118 |
| Mayo | TCX | TCXturquoise | TCXturquoise_1 | 251 |
| Mayo | TCX | TCXturquoise | TCXturquoise_2 | 220 |
| Mayo | TCX | TCXturquoise | TCXturquoise_3 | 171 |
| Mayo | TCX | TCXyellow | TCXyellow_1 | 975 |
| Mayo | TCX | TCXyellow | TCXyellow_2 | 189 |
| MSBB | FP | FPblue | FPblue_1 | 558 |
| MSBB | FP | FPblue | FPblue_2 | 229 |
| MSBB | FP | FPblue | FPblue_3 | 105 |
| MSBB | FP | FPblue | FPblue_4 | 101 |
| MSBB | FP | FPbrown | FPbrown_1 | 258 |
| MSBB | FP | FPbrown | FPbrown_2 | 99 |
| MSBB | FP | FPturquoise | FPturquoise_1 | 393 |
| MSBB | FP | FPturquoise | FPturquoise_2 | 100 |
| MSBB | FP | FPyellow | FPyellow_1 | 1328 |
| MSBB | FP | FPyellow | FPyellow_2 | 226 |
| MSBB | FP | FPyellow | FPyellow_3 | 145 |
| MSBB | FP | FPyellow | FPyellow_4 | 101 |
| MSBB | FP | FPyellow | FPyellow_5 | 101 |
| MSBB | IFG | IFGblue | IFGblue_1 | 541 |
| MSBB | IFG | IFGblue | IFGblue_2 | 121 |
| MSBB | IFG | IFGblue | IFGblue_3 | 100 |
| MSBB | IFG | IFGbrown | IFGbrown_1 | 1415 |
| MSBB | IFG | IFGbrown | IFGbrown_2 | 187 |
| MSBB | IFG | IFGbrown | IFGbrown_3 | 163 |
| MSBB | IFG | IFGbrown | IFGbrown_4 | 140 |
| MSBB | IFG | IFGturquoise | IFGturquoise_1 | 346 |
| MSBB | IFG | IFGturquoise | IFGturquoise_2 | 139 |
| MSBB | IFG | IFGturquoise | IFGturquoise_3 | 127 |
| MSBB | IFG | IFGyellow | IFGyellow_1 | 277 |
| MSBB | PHG | PHGblue | PHGblue_1 | 152 |
| MSBB | PHG | PHGblue | PHGblue_2 | 126 |

| | | | | |
|---|---|---|---|---|
| MSBB | PHG | PHGblue | PHGblue_3 | 124 |
| MSBB | PHG | PHGblue | PHGblue_4 | 113 |
| MSBB | PHG | PHGbrown | PHGbrown_1 | 1144 |
| MSBB | PHG | PHGbrown | PHGbrown_2 | 194 |
| MSBB | PHG | PHGbrown | PHGbrown_3 | 117 |
| MSBB | PHG | PHGbrown | PHGbrown_4 | 101 |
| MSBB | PHG | PHGgreen | PHGgreen_1 | 578 |
| MSBB | PHG | PHGturquoise | PHGturquoise_1 | 399 |
| MSBB | PHG | PHGturquoise | PHGturquoise_2 | 77 |
| MSBB | PHG | PHGyellow | PHGyellow_1 | 335 |
| MSBB | STG | STGblue | STGblue_1 | 287 |
| MSBB | STG | STGblue | STGblue_2 | 234 |
| MSBB | STG | STGblue | STGblue_3 | 148 |
| MSBB | STG | STGbrown | STGbrown_1 | 1088 |
| MSBB | STG | STGbrown | STGbrown_2 | 158 |
| MSBB | STG | STGbrown | STGbrown_3 | 157 |
| MSBB | STG | STGbrown | STGbrown_4 | 122 |
| MSBB | STG | STGturquoise | STGturquoise_1 | 103 |
| MSBB | STG | STGyellow | STGyellow_1 | 645 |
| ROSMAP | DLPFC | DLPFCblue | DLPFCblue_1 | 452 |
| ROSMAP | DLPFC | DLPFCblue | DLPFCblue_2 | 242 |
| ROSMAP | DLPFC | DLPFCblue | DLPFCblue_3 | 183 |
| ROSMAP | DLPFC | DLPFCblue | DLPFCblue_4 | 175 |
| ROSMAP | DLPFC | DLPFCbrown | DLPFCbrown_1 | 182 |
| ROSMAP | DLPFC | DLPFCbrown | DLPFCbrown_2 | 117 |
| ROSMAP | DLPFC | DLPFCturquoise | DLPFCturquoise_1 | 460 |
| ROSMAP | DLPFC | DLPFCturquoise | DLPFCturquoise_2 | 301 |
| ROSMAP | DLPFC | DLPFCyellow | DLPFCyellow_1 | 1296 |
| ROSMAP | DLPFC | DLPFCyellow | DLPFCyellow_2 | 163 |
| ROSMAP | DLPFC | DLPFCyellow | DLPFCyellow_3 | 105 |

**Supplemental Table 4: Summary of Submodules**
Submodules were generated from existing modules generated for each tissue region. The number of genes in each submodule is reported. 68 submodules were generated for this study.

| Cohort | Brain Region | Subtype | Cases |
|---|---|---|---|
| Mayo | TCX | A | 23 |
| Mayo | TCX | B | 37 |
| Mayo | TCX | C | 20 |
| Mayo | TCX | Control | 182 |
| MSBB | FP | A | 39 |
| MSBB | FP | B | 51 |
| MSBB | FP | Control | 124 |
| MSBB | PHG | A | 43 |
| MSBB | PHG | B | 21 |
| MSBB | PHG | Control | 96 |
| ROSMAP | DLPFC | A | 80 |
| ROSMAP | DLPFC | B | 72 |
| ROSMAP | DLPFC | Control | 471 |

**Supplemental Table 5: LOAD Case Subtypes for Selected Brain Regions**
Subtypes were generated for the TCX, FP, PHG, and DLPFC regions. 3 clusters were generated for TCX and 2 clusters were generated for the rest. The number of cases in each subtype are reported.

**Supplemental Table 6: GO Term Annotations of Submodules**

Enrichment of genes in each submodule was assessed using the clusterProfiler R package for GO terms. GO terms for each submodule are reported (see attached Excel workbook).

**Supplemental Table 7: KEGG Pathway Annotations of Submodules**

Enrichment of genes in each submodule was assessed using the clusterProfiler R package for KEGG pathways. KEGG pathways for each submodule are reported (see attached Excel workbook).

**Supplemental Table 8: Reactome Pathway Annotations of Submodules**

Enrichment of genes in each submodule was assessed using the ReactomePA R package for Reactome pathways. Reactome pathways for each submodule are reported (see attached Excel workbook).

**Supplemental Table 9: Significant SNP Associations from TCX Region Analyses**

Significant SNPs that were associated at a genome-wide suggestive level with either a submodule eigengene or the subtype specificity metric are reported. RefSNP IDs are provided if available for the positions, which are aligned to the hg19 human genome build (see attached Excel workbook).

**Supplemental Table 10: Significant SNP Associations from PHG Region Analyses**

Significant SNPs that were associated at a genome-wide suggestive level with either a submodule eigengene or the subtype specificity metric are reported. RefSNP IDs are provided if available for the positions, which are aligned to the hg19 human genome build (see attached Excel workbook).

**Supplemental Table 11: Significant SNP Associations from FP Region Analyses**

Significant SNPs that were associated at a genome-wide suggestive level with either a submodule eigengene or the subtype specificity metric are reported. RefSNP IDs are provided if available for the positions, which are aligned to the hg19 human genome build (see attached Excel workbook).

**Supplemental Table 12: Significant SNP Associations from DLPFC Region Analyses**

Significant SNPs that were associated at a genome-wide suggestive level with either a submodule eigengene or the subtype specificity metric are reported. RefSNP IDs are provided if available for the positions, which are aligned to the hg19 human genome build (see attached Excel workbook).

**Supplemental Table 13: KEGG Pathway Annotations of Differentially Expressed Genes in ROSMAP Subtypes**

Differentially expressed genes between controls and each subtype were enriched for KEGG pathway annotations using the clusterProfiler R package. KEGG pathways and associated scores are reported (see attached Excel workbook).

**Supplemental Table 14: Reactome Pathway Annotations of Differentially Expressed Genes in ROSMAP Subtypes**
Differentially expressed genes between controls and each subtype were enriched for Reactome pathway annotations using the ReactomePA R package. Reactome pathways and associated scores are reported (see attached Excel workbook).

**Supplemental Table 15: Genome-Wide Suggestive SNPs in DLPFC Replicated in TCX**
SNPs that were found to be genome-wide suggestive in the DLPFC analyses were assessed for replication in the analyses run for the TCX region. A p-value cutoff of 0.05 was used for the TCX region. The analysis that generated the highest p-value for the SNP in DLPFC and TCX are reported, along with the p-values from each (see attached Excel workbook).

**Supplemental Table 16: Genome-Wide Suggestive SNPs in DLPFC Replicated in FP**
SNPs that were found to be genome-wide suggestive in the DLPFC analyses were assessed for replication in the analyses run for the FP region. A p-value cutoff of 0.05 was used for the FP region. The analysis that generated the highest p-value for the SNP in DLPFC and FP are reported, along with the p-values from each (see attached Excel workbook).

**Supplemental Table 17: Genome-Wide Suggestive SNPs in DLPFC Replicated in PHG**
SNPs that were found to be genome-wide suggestive in the DLPFC analyses were assessed for replication in the analyses run for the PHG region. A p-value cutoff of 0.05 was used for the PHG region. The analysis that generated the highest p-value for the SNP in DLPFC and PHG are reported, along with the p-values from each (see attached Excel workbook).

**Supplementary Table 18: Genome-Wide Suggestive SNPs in DLPFC Replicated in the NHGRI-EBI Catalog**
Genome-wide suggestive SNPs from the DLPFC region were assessed for replication in the summary SNPs provided by the NHGRI-EBI catalog (see attached Excel workbook).
